# Scientific Article Dietary electrolyte balance on serum hematological and biochemical parameters of broilers under heat stress

**DOI:** 10.1101/2021.01.11.426185

**Authors:** Danilo Rodrigo Silva e Silva, Francinete Alves de Sousa Moura, Vânia Batista de Sousa Lima, Patrícia Miranda Lopes, Janaina de Fatima Saraiva Cardoso, Alan Costa do Prado, Luciana Pereira Machado, Richard Átila de Sousa, Daniel Biagiotti, Leilane Rocha Barros Dourado

**Affiliations:** Campus Petrônio Portela, Federal University of Piauí, Teresina, Piauí, Brazil; Campus Teacher Cinobelina Elvas, Federal University of Piauí, Bom Jesus, Piauí, Brazil; Department of Veterinary Medicine, Campus Sertão, Federal University of Rio Grande do Sul, Sertão, Rio Grande do Sul, Brazil; Department of Veterinary Medicine, Federal University of Fronteira Sul, Chapecó, Santa Catarina, Brazil

## Abstract

The aim of this study was to evaluate the effect of different dietary electrolyte balance (DEB) levels over the hematological parameters and the biochemical profile of broilers under stress caused by cyclic heat. We used 1050 male broilers from Ross lineage, for two experimental phases: phase 1 (broilers from 1 until 21 days old) and phase 2 (broilers from 22 until 42 days old). The broilers were distributed under a completely randomized design, built for five treatments (110; 175; 240; 305; 370 mEq of DEB/kg of ration) and seven repetitions. For the hematological analyses were used two broilers in each repetition, one to blood count evaluation (Hemogram) and another one to the biochemistry evaluation, collected in the end of each experimental phase. During the phase of 1 - 21 days old, the variable hemoglobin number (Heme), mean corpuscular volume (MCV) and Serum Chloride (Cl) presented a linear effect (p<0,05) DEB levels. There was no effects (p>0,05) over the corpuscular volume (CV), total plasma protein (TPP), leukocytes number (Leu), heterophiles (Het), lymphocytes (Lyn), eosinophils (Eos), basophils (Bas), monocytes (Mon), calcium (Ca), phosphor (P), uric acid (UA) e urea (U). During the phase two of 22 – 42 days old there was effect (p<0,05) quadratic to the variables Leu and Het and linear to the variable P and significative difference (p<0,05) between the variable means UA at the levels of DEB tested.

There was no effect (p>0,05) over the variable CV, Heme, MCV, TPP, Lyn, Eos, Bas, Mon, Cl, Ca and U on the DEB levels tested. The supplementation of the DEB levels 110, 175 e 240, 305 and 370 to the phase from 1 until 21 days old presented great results in the hematological evaluation and biochemical, because it provided possibly a longer useful life to the Hem without promoting a metabolic imbalance. In the phase from 22 until 42 days old the level of 268 to 280 mEq of DEB/kg of ration presented a better answer from the broilers immune systemic.

## Introduction

One of the biggest obstacles about broilers creation is the environment in which they grow up, mainly in tropical and subtropical regions where predominate high temperatures, in which the chronic heat stress (over 30 °C) can harm poultry production [1]. As soon as the ambient temperature exceeds the thermal comfort temperature of the broiler, where the metabolic rate is minimal and the homeotherm is maintained with less energy expenditure, the heat stress begins [2]. Consequently, the broilers can suffer metabolic imbalance (acidosis or alkalosis), decreased feed consumption and low growth rate, impaired health and increased mortality [3].

Respiratory alkalosis occurs in the heat stress situation due to an increase in the respiratory rate, resulting in excessive carbon dioxide losses (CO_2_). Thus, the partial pressure of CO_2_ (pCO_2_) decreases, leading to a drop in the concentration of carbonic acid (H_2_CO_3_) and hydrogen (H^+^). On the other hand, as the stress intensifies, there is an increase in the partial pressure of (pCO_2_) with an increase in the level of H^+^, causing respiratory acidosis [4].

In tropical production systems, several strategies are used to mitigate the effects of heat stress, such as modifying the environment through the projection of acclimatized facilities, managements such as feeding and temperature control of drinking water, in addition to changes in the diet composition in order to promote greater intake or compensate for low food consumption with or without the use of additives such as vitamins and minerals [5].

Dietary electrolyte supplementation has been used to maintain the acid-base balance in the blood, minimizing the deleterious effects of heat stress [6]. Mongin [7] and Johnson and Karunajeewa [8] were the first ones to relate the applicability of the electrolytes use, with the intent of balance their lose from the broilers under heat stress, these authors observed better performance in broilers fed with diets under dietary electrolyte balance (DEB) 250 and 180 to 300 mEq / kg, respectively. Ahmad et al [6] found out that a DEB between 50 and 250 mEq / kg promotes acidobasic normalization of broilers under heat stress, however, DEB of 0 mEq / kg causes growth depression due to lower feed consumption, because of blood acidosis, and the DEB 350 mEq / kg, causes alkalosis, associated with a high level of NaHCO_3_ [6].

In the hematological and biochemical evaluation, it has already been observed that heat stress causes an increase in hematocrit (justified by the effect of dehydration caused by vomiting and diarrhea) and in the number of red blood cells through an increase in erythropoiesis [9]. The H/L ratio rises as a consequence of the heterophile’s increase and reduction of lymphocyte, this relationship is a sensitive index of chronic stress in broilers [11]. There has also been observed a decrease in serum total plasma proteins and uric acid, an increase and/or decrease in chlorine, among other changes [12, 13].

Therefore, the evaluation of the efficiency of the inclusion of DEB can be carried out through blood parameters, since these parameters are very susceptible to temperature variation, constituting into an important indication of physiological responses to stress [11]. Based on these evidences, the effect of the electrolyte balance (110; 175; 240; 305; 370 mEq/kg) on the hemogram and the serum biochemical profile of broilers will be evaluated, in order to identify the efficiency of these diets in situations of cyclic heat stress.

## Materials and methods

This study was conducted in strict accordance with the recommendations of the guide for care and use of laboratory animals from the National Institutes of Health. The protocol was approved by the Ethics Committee on Animal Experimentation at the Federal University of Piauí (Piauí, Brazil) under the number 075/15.

### Birds and diets

The experiment was conducted in the poultry sector of the Federal University of Piauí, located in Bom Jesus-PI, Brazil (latitude: 9° 4′30″ south and longitude: 44° 21′26″ west). 1050 male Ross broilers were used for two experimental phases: phase 1 (broilers from 1 to 21 days old) and phase 2 (broilers from 22 to 42 days old), 525 in each phase. The animals were distributed in a completely randomized design, consisting of five treatments (levels of electrolyte balance) and seven replications of 15 broilers, and for hematological analysis two broilers from each repetition were used, one for hemogram evaluation and another for biochemical evaluation collected at the end of each experimental phase, 72 per experimental phase, totaling 144 birds.

The experimental diets were formulated to attend the nutritional requirements of the broilers in each phase (1 to 21 and 22 to 42 days), considering the requirements and chemical composition of the ingredients, as described by Rostagno et al. [14], except for sodium, chlorine and potassium that were adjusted to define the electrolyte balances tested, and also to assess the levels of these elements in the ingredients used (Table 1, 2, 3 and 4). To obtain the DEB levels (110; 175; 240; 305; 370 mEq / kg), ammonia chloride (NH_4_Cl), potassium carbonate (K_2_CO_3_) and/or sodium bicarbonate (NaHCO_3_) were added to the diets, replacing the inert material. Water and feed were provided *ad libitum* to the animals.

**Table 1.** Centesimal composition of experimental diets for broilers from 1 to 7 days old.

| Ingredients | Dietary electrolyte balance – mEq/kg |  |  |  |  |
| --- | --- | --- | --- | --- | --- |
|  | 110 | 175 | 240 | 305 | 370 |
| Grain corn | 54,283 | 54,283 | 54,283 | 54,283 | 54,283 |
| Soybean meal | 35,409 | 35,409 | 35,409 | 35,409 | 35,409 |
| Soy oil | 3,373 | 3,373 | 3,373 | 3,373 | 3,373 |
| Dicalcium phosphate | 1,862 | 1,862 | 1,862 | 1,862 | 1,862 |
| Calcitic limestone | 0,905 | 0,905 | 0,905 | 0,905 | 0,905 |
| DL-methionine | 0,221 | 0,221 | 0,221 | 0,221 | 0,221 |
| L- lysine | 0,601 | 0,601 | 0,601 | 0,601 | 0,601 |
| L- threonine | 0,165 | 0,165 | 0,165 | 0,165 | 0,165 |
| L- arginine | 0,133 | 0,133 | 0,133 | 0,133 | 0,133 |
| L-valine | 0,150 | 0,150 | 0,150 | 0,150 | 0,150 |
| Common salt | 0,540 | 0,540 | 0,540 | 0,540 | 0,540 |
| Suppl. Min. Vit.<sup>1</sup> | 1,000 | 1,000 | 1,000 | 1,000 | 1,000 |
| Inert | 0,920 | 1,267 | 1,007 | 0,507 | - |
| NaHCO<sub>3</sub> | - | - | 0,173 | 0,487 | 0,840 |
| K<sub>2</sub>CO<sub>3</sub> | - | - | 0,180 | 0,367 | 0,520 |
| NH<sub>4</sub>CL | 0,440 | 0,093 | - | - | - |
| Total | 100,000 | 100,000 | 100,000 | 100,000 | 100,000 |
| Calculated Composition |  |  |  |  |  |
| Linoleic acid (%) | 3,095 | 3,095 | 3,095 | 3,095 | 3,095 |
| Calcium (%) | 0,971 | 0,971 | 0,971 | 0,971 | 0,971 |
| Energ. Met. Poultry (Mcal / kg) | 2,975 | 2,975 | 2,975 | 2,975 | 2,975 |
| Available phosphorus (%) | 0,463 | 0,463 | 0,463 | 0,463 | 0,463 |
| Lysine dig. Birds (%) | 1,307 | 1,307 | 1,307 | 1,307 | 1,307 |
| Met. + Cyst dig. Birds (%) | 0,967 | 0,967 | 0,967 | 0,967 | 0,967 |
| Dig Arginine. Birds (%) | 1,398 | 1,398 | 1,398 | 1,398 | 1,398 |
| Threonine dig. Birds (%) | 0,863 | 0,863 | 0,863 | 0,863 | 0,863 |
| Dig tryptophan Birds (%) | 0,235 | 0,235 | 0,235 | 0,235 | 0,235 |
| Valina dig. Birds (%) | 1,006 | 1,006 | 1,006 | 1,006 | 1,006 |
| Crude protein (%) | 21,173 | 21,173 | 21,173 | 21,173 | 21,173 |
| Potassium (%) | 1,116 | 1,116 | 1,216 | 1,324 | 1,411 |
| Sodium (%) | 0,225 | 0,225 | 0,273 | 0,358 | 0,455 |
| Chlorine (%) | 0,962 | 0,736 | 0,674 | 0,674 | 0,674 |

<sup>1</sup>A supplies/kg of diet. Pre-initial: Folic acid - 200.00mg; Biotin - 10.00mg; Chlorohydroxyquinoline - 7500.00mg; Zn - 17.50g; Vit. A - 1680000,00UI; Vit. B1 - 436.50mg; Vit. 2400.00mg B12; Vit. B2 - 1200.00mg; Vit. B6 - 624mg; Vit. D3 - 400000,00UI; Vit. E - 3500.00UI; Vit. K3 - 360.00mg; Niacin - 8399.00mg; Nicarbazine - 25.00g; Pantothenic acid - 3120.00mg; Choline - 78.10g; Se - 75.00mg; Fe 11.25g; Mn - 18.74g; Cu - 1997.00mg; I - 187.00mg.

**Table 2.**
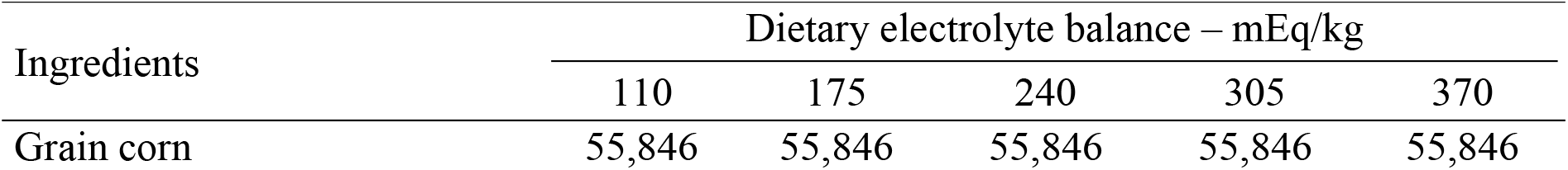
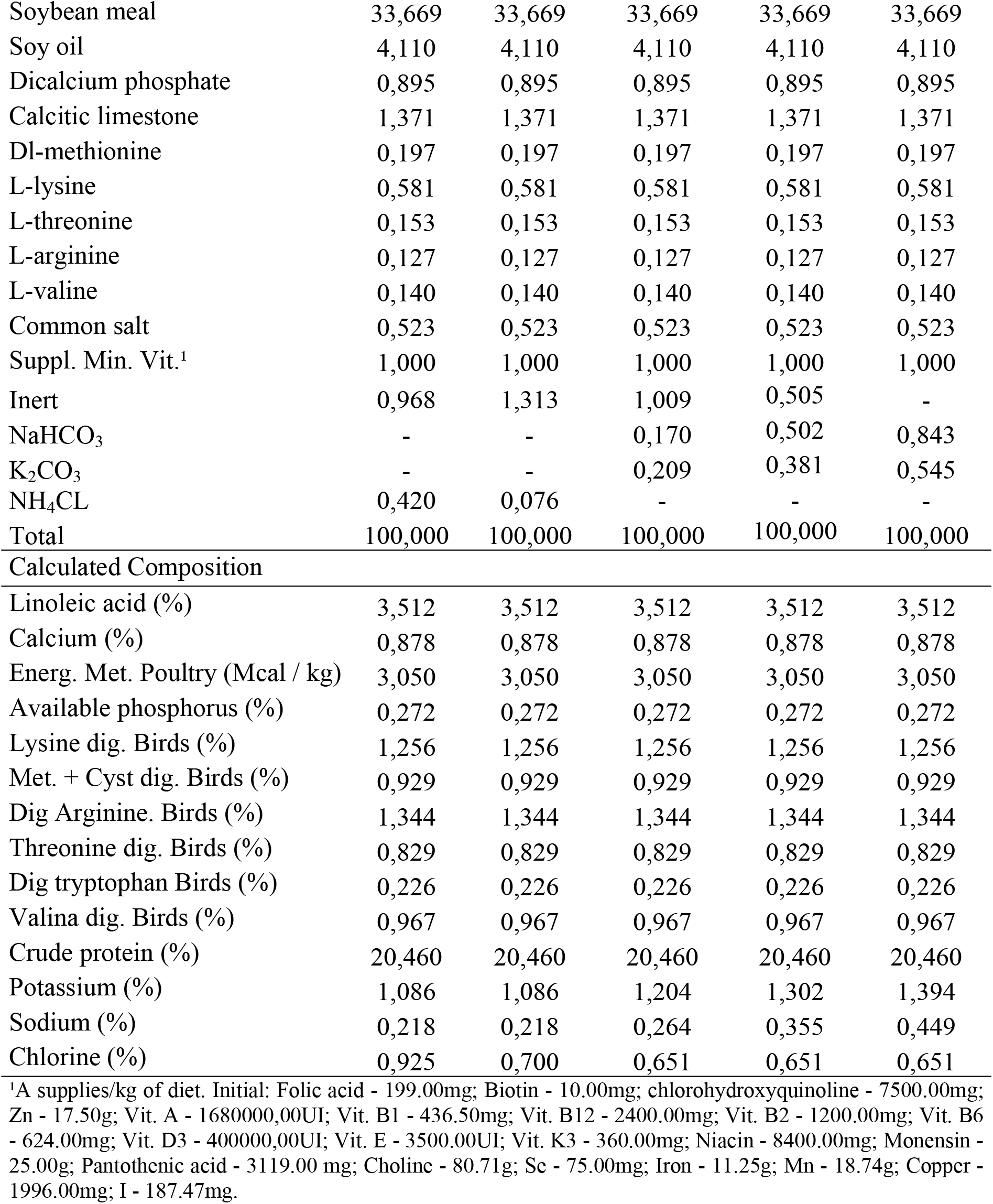
Centesimal composition of experimental diets for broilers from 8 to 21 days old.

**Table 3.** Centesimal composition of experimental diets for broilers from 22 to 33 days old.

| Ingredients | Dietary electrolyte balance – mEq/kg |  |  |  |  |
| --- | --- | --- | --- | --- | --- |
|  | 110 | 175 | 240 | 305 | 370 |
| Grain corn | 59,745 | 59,745 | 59,745 | 59,745 | 59,745 |
| Soybean meal | 29,055 | 29,055 | 29,055 | 29,055 | 29,055 |
| Soy oil | 4,970 | 4,970 | 4,970 | 4,970 | 4,970 |
| Dicalcium phosphate | 1,451 | 1,451 | 1,451 | 1,451 | 1,451 |
| Calcitic limestone | 0,686 | 0,686 | 0,686 | 0,686 | 0,686 |
| Dl-methionine | 0,236 | 0,236 | 0,236 | 0,236 | 0,236 |
| L-lysine | 0,537 | 0,537 | 0,537 | 0,537 | 0,537 |
| L-threonine | 0,125 | 0,125 | 0,125 | 0,125 | 0,125 |
| L-arginine | 0,114 | 0,114 | 0,114 | 0,114 | 0,114 |
| L-valine | 0,115 | 0,115 | 0,115 | 0,115 | 0,115 |
| Common salt | 0,500 | 0,500 | 0,500 | 0,500 | 0,500 |
| Suppl. Min. Vit.<sup>1</sup> | 1,000 | 1,000 | 1,000 | 1,000 | 1,000 |
| Inert | 1,109 | 1,455 | 1,003 | 0,503 | - |
| NaHCO<sub>3</sub> | - | - | 0,183 | 0,490 | 0,819 |
| K<sub>2</sub>CO<sub>3</sub> | - | - | 0,280 | 0,473 | 0,647 |
| NH<sub>4</sub>CL | 0,357 | 0,011 | - | - | - |
| Total | 100,000 | 100,000 | 100,000 | 100,000 | 100,000 |
| Calculated Composition |  |  |  |  |  |
| Linoleic acid (%) | 4,019 | 4,019 | 4,019 | 4,019 | 4,019 |
| Calcium (%) | 0,758 | 0,758 | 0,758 | 0,758 | 0,758 |
| Energ. Met. Poultry (Mcal / kg) | 3,150 | 3,150 | 3,150 | 3,150 | 3,150 |
| Available phosphorus (%) | 0,374 | 0,374 | 0,374 | 0,374 | 0,374 |
| Lysine dig. Birds (%) | 1,124 | 1,124 | 1,124 | 1,124 | 1,124 |
| Met. + Cyst dig. Birds (%) | 0,832 | 0,832 | 0,832 | 0,832 | 0,832 |
| Dig Arginine. Birds (%) | 1,203 | 1,203 | 1,203 | 1,203 | 1,203 |
| Threonine dig. Birds (%) | 0,742 | 0,742 | 0,742 | 0,742 | 0,742 |
| Dig tryptophan Birds (%) | 0,202 | 0,202 | 0,202 | 0,202 | 0,202 |
| Valina dig. Birds (%) | 0,865 | 0,865 | 0,865 | 0,865 | 0,865 |
| Crude protein (%) | 18,621 | 18,621 | 18,621 | 18,621 | 18,621 |
| Potassium (%) | 1,003 | 1,003 | 1,161 | 1,271 | 1,369 |
| Sodium (%) | 0,208 | 0,208 | 0,258 | 0,342 | 0,432 |
| Chlorine (%) | 0,834 | 0,609 | 0,602 | 0,602 | 0,602 |

<sup>1</sup>A supplies/kg of diet. Growth: Folic acid - 162.50mg; Chlorohydroxyquinoline - 7500.00mg; Zn - 17.50g; Vit. A - 1400062,50UI; Vit. B1 - 388.00mg; Vit. B12 - 2000.00mcg; Vit. B2 - 1000.00 Mg; Vit. B6 - 520.00mg; Vit. D3 - 360012,00UI; Vit. E - 2500.00UI; Vit. K3 - 300.00mg; Niacin - 7000.00mg; Salinomycin - 16.50g; Pantothenic acid - 2600.00mg; Choline - 71.59g; Se - 75.00mg; Fe - 11.25g; Mn - 18.74g; Cu - 1996.00mg; I - 187.47mg.

**Table 4.**
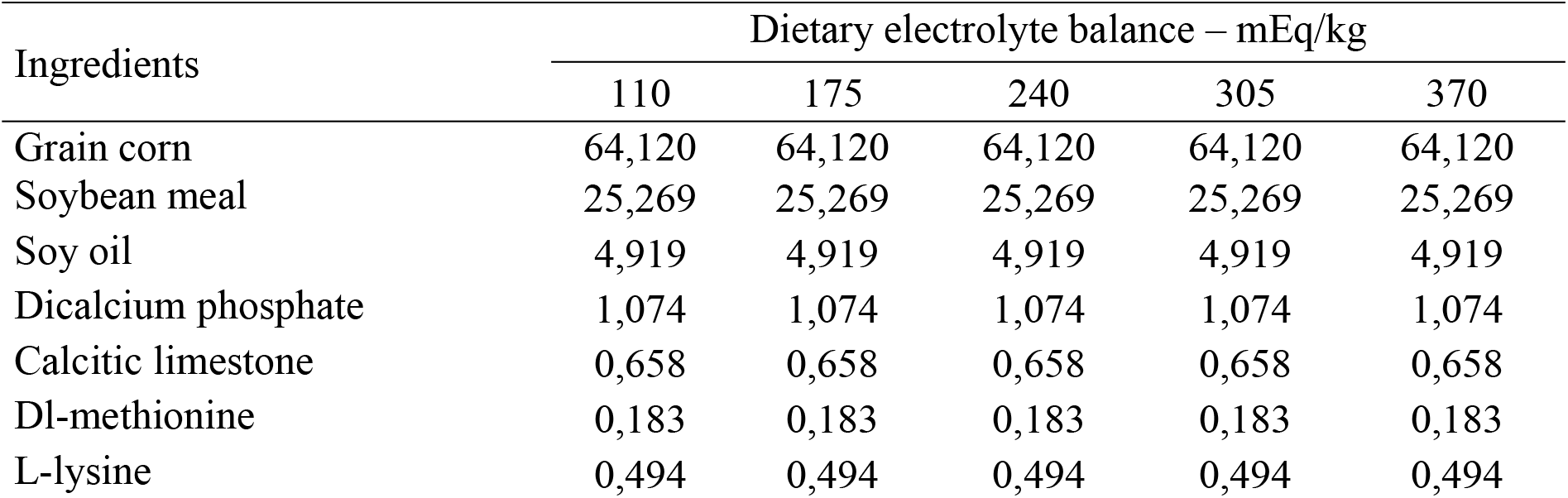
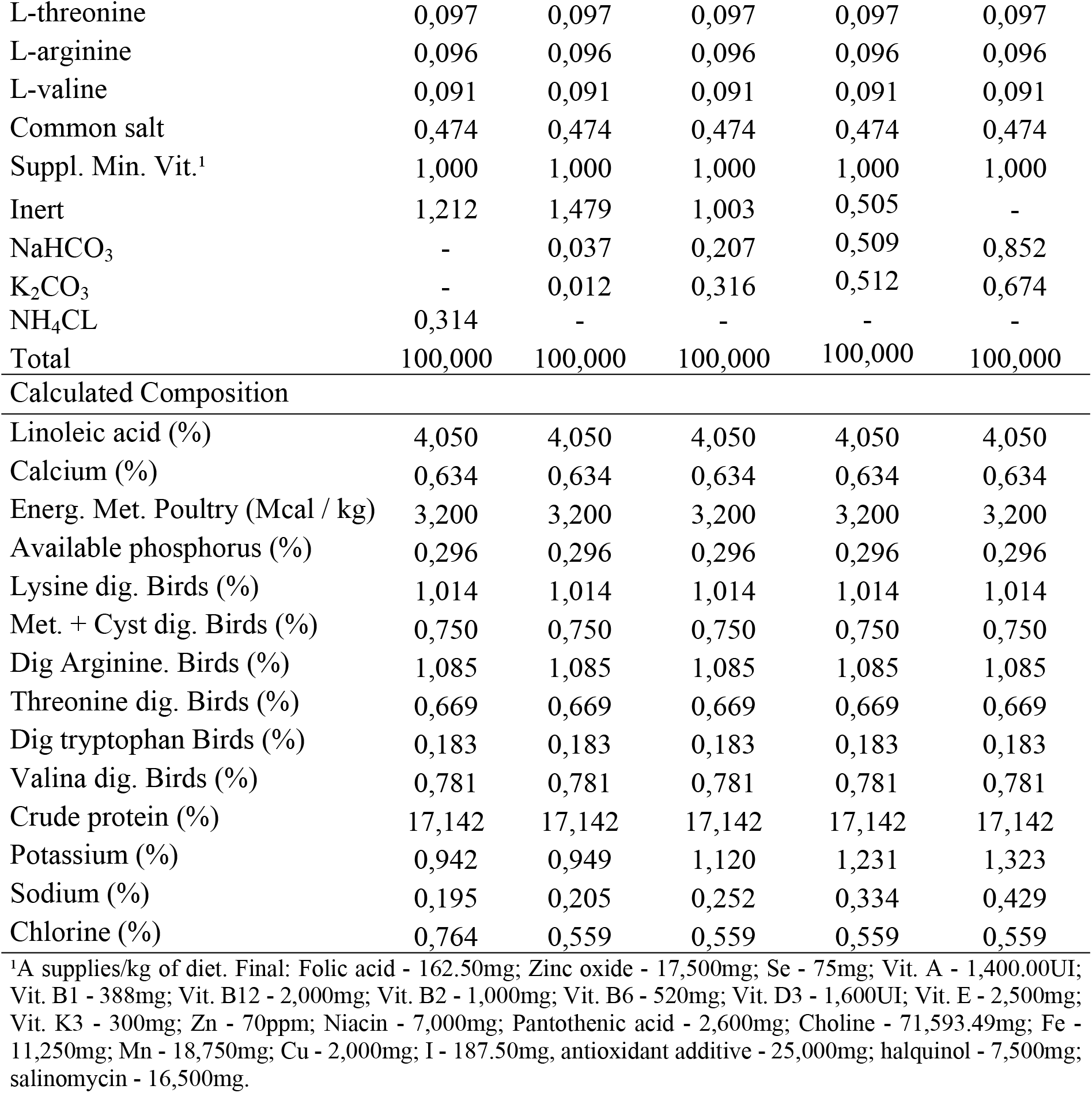
Centesimal composition of experimental diets for broilers from 34 to 42 days old

| Ingredients | Dietary electrolyte balance – mEq/kg |  |  |  |  |
| --- | --- | --- | --- | --- | --- |
|  | 110 | 175 | 240 | 305 | 370 |
| Grain corn | 64,120 | 64,120 | 64,120 | 64,120 | 64,120 |
| Soybean meal | 25,269 | 25,269 | 25,269 | 25,269 | 25,269 |
| Soy oil | 4,919 | 4,919 | 4,919 | 4,919 | 4,919 |
| Dicalcium phosphate | 1,074 | 1,074 | 1,074 | 1,074 | 1,074 |
| Calcitic limestone | 0,658 | 0,658 | 0,658 | 0,658 | 0,658 |
| Dl-methionine | 0,183 | 0,183 | 0,183 | 0,183 | 0,183 |
| L-lysine | 0,494 | 0,494 | 0,494 | 0,494 | 0,494 |
| L-threonine | 0,097 | 0,097 | 0,097 | 0,097 | 0,097 |
| L-arginine | 0,096 | 0,096 | 0,096 | 0,096 | 0,096 |
| L-valine | 0,091 | 0,091 | 0,091 | 0,091 | 0,091 |
| Common salt | 0,474 | 0,474 | 0,474 | 0,474 | 0,474 |
| Suppl. Min. Vit.<sup>1</sup> | 1,000 | 1,000 | 1,000 | 1,000 | 1,000 |
| Inert | 1,212 | 1,479 | 1,003 | 0,505 | - |
| NaHCO<sub>3</sub> | - | 0,037 | 0,207 | 0,509 | 0,852 |
| K<sub>2</sub>CO<sub>3</sub> | - | 0,012 | 0,316 | 0,512 | 0,674 |
| NH<sub>4</sub>CL | 0,314 | - | - | - | - |
| Total | 100,000 | 100,000 | 100,000 | 100,000 | 100,000 |
| Calculated Composition |  |  |  |  |  |
| Linoleic acid (%) | 4,050 | 4,050 | 4,050 | 4,050 | 4,050 |
| Calcium (%) | 0,634 | 0,634 | 0,634 | 0,634 | 0,634 |
| Energ. Met. Poultry (Mcal / kg) | 3,200 | 3,200 | 3,200 | 3,200 | 3,200 |
| Available phosphorus (%) | 0,296 | 0,296 | 0,296 | 0,296 | 0,296 |
| Lysine dig. Birds (%) | 1,014 | 1,014 | 1,014 | 1,014 | 1,014 |
| Met. + Cyst dig. Birds (%) | 0,750 | 0,750 | 0,750 | 0,750 | 0,750 |
| Dig Arginine. Birds (%) | 1,085 | 1,085 | 1,085 | 1,085 | 1,085 |
| Threonine dig. Birds (%) | 0,669 | 0,669 | 0,669 | 0,669 | 0,669 |
| Dig tryptophan Birds (%) | 0,183 | 0,183 | 0,183 | 0,183 | 0,183 |
| Valina dig. Birds (%) | 0,781 | 0,781 | 0,781 | 0,781 | 0,781 |
| Crude protein (%) | 17,142 | 17,142 | 17,142 | 17,142 | 17,142 |
| Potassium (%) | 0,942 | 0,949 | 1,120 | 1,231 | 1,323 |
| Sodium (%) | 0,195 | 0,205 | 0,252 | 0,334 | 0,429 |
| Chlorine (%) | 0,764 | 0,559 | 0,559 | 0,559 | 0,559 |

<sup>1</sup>A supplies/kg of diet. Final: Folic acid - 162.50mg; Zinc oxide - 17,500mg; Se - 75mg; Vit. A - 1,400.00UI; Vit. B1 - 388mg; Vit. B12 - 2,000mg; Vit. B2 - 1,000mg; Vit. B6 - 520mg; Vit. D3 - 1,600UI; Vit. E - 2,500mg; Vit. K3 - 300mg; Zn - 70ppm; Niacin - 7,000mg; Pantothenic acid - 2,600mg; Choline - 71,593.49mg; Fe - 11,250mg; Mn - 18,750mg; Cu - 2,000mg; I - 187.50mg, antioxidant additive - 25,000mg; halquinol - 7,500mg; salinomycin - 16,500mg.

Throughout the experimental period, a continuous light program (24 hours of natural + artificial light) was adopted, using 60 Watt incandescent lamps, applying thermal stress to these animals, considering that the temperature in Bom Jesus tends to go beyond the thermal comfort zone for broilers, which according to Abreu and Abreu [15] is around 26 to 29 ° C for broilers with 21 days and 20°C for broilers with 42 days, with relative humidity between 60 to 70% for both phases. The temperature and relative humidity of the air inside the shed was measured daily, throughout the experimental period, by thermohygrometers.

### Sample collection

For laboratory analysis of the blood count and serum biochemistry, 0.5 to 3 mL of venous blood were collected, depending on the size of the bird and not exceeding 1% of the animal’s weight, by puncture of the jugular vein, medial metatarsal or ulnar vein, in syringes containing 10% EDTA anticoagulant (ethylenediaminetetraacetic acid) for blood count and syringes without anticoagulant for biochemical analysis [16]. Blood samples were taken in the two experimental phases, the first phase was performed on the 20th day of age and the second phase on the 41st day of age.

### Hemogram

Immediately after collection, blood smears were made, subsequently stained with hematological dye type Romanowsky (Panotic Rapid®) and displacement under an optical microscope, without a 1000x increase, for differential assessment of leukocytes. Then the absolute red blood cell and leukocyte counts were performed in a Neubauer camera using 0.01% toluidine blue. Hematocrit was also determined by centrifuging a blood sample in a microhematocrit capillary and calculating the mean corpuscular volume (MCV) according to Hendrix [17].

### Biochemistry

To evaluate serum biochemistry, an automatic biochemical analyzer (Automatic Analyzer Chem Well T - Labtest, Lagoa Santa/MG, Brazil) from Hospital Veterinário Universitário, Dr. Jeremias Pereira da Silva, from UFPI, Teresina / PI, Brazil, was used, using commercial kits. The serum concentrations of urea (UV enzyme methodology), calcium (colorimetric methodology), phosphorus (Daly and modified Ertingshausen methodology), chloride (Mercury Thiocyanate methodology) and the serum uric acid concentration (enzymatic colorimetric Trinder method) were determined. All biochemical reactions were processed as directed by the manufacturers.

### Total plasma protein (TPP)

The determination of TPP was performed after centrifuging blood in a microhematocrit capillary and reading the plasma protein concentration by refractometer [16].

### Statistical analysis

The data were subjected to a normality test (Cramer-Von Mises), with analysis of variance (ANOVA) by the GLM procedure of the *Statistical Analysis System - SAS software, version* The SNK test with α = 0.05 probability was used to compare means. Estimates of dietary electrolyte balance were defined using polynomial regression.

## Results

### Hemogram and Toral Plasma Protein

The hemogram results are presented on tables 5 and 6 to the phase from 1 – 21 days old and tables 7 and 8 to the phase from 21 - 42 days old. For the 1-21 days old phase, the number of red blood cells (Heme) and MCV variables showed a linear effect (p <0.05) because the electrolyte balance levels. There was no effect (p> 0.05) on the variables globular volume (GV), TPP, number of leukocytes (Leu), heterophiles (Het), lymphocytes (Lyn), eosinophils (Eos), basophils (Bas), monocytes (Mon) and the H / L ratio as a function of the DEB levels tested. A value above the reference of the species was observed for the variable Het at levels 175, 240 and 370 mEq / kg and for variable Mon in all tested levels. The average maximum and minimum temperature during this phase were 34.2 and 21.4 °C, respectively, and the maximum and minimum humidity were 62.57 and 29.43%, and for the day of collection the maximum and minimum temperature were respectively 34.2 and 20.7 °C, and the maximum and minimum humidity 61 and 28%, characterizing chronic stress by cyclic heat.

**Table 5.** Levels of dietary electrolyte balance on the erythrogram and total plasma proteins (TPP) of broilers from 01 to 21 days of age.

| DEB (mEq/Kg) | Heme ($\times 10^6/\mu\text{L}$) | GV (%) | MCV (fL) | TPP (g/dL) |
| --- | --- | --- | --- | --- |
| 110 | 1,81 | 29,0 | 155,4 | 3,4 |
| 175 | 1,81 | 29,8 | 163,1 | 3,3 |
| 240 | 2,06 | 28,3 | 147,8 | 3,4 |
| 305 | 1,98 | 29,4 | 143,8 | 3,3 |
| 370 | 2,08 | 30,3 | 145,9 | 3,5 |
| RV | 1,98 ± 0,28<sup>”</sup> | 22 – 35<sup>*</sup> | 170,48 ± 24,83<sup>”</sup> | 2,5 - 4,5<sup>**</sup> |
| Probability | 0,1342 | 0,2189 | 0,1085 | 0,5222 |
| Regression | L<sup>1</sup> | Ns | L<sup>2</sup> | Ns |
| CV (%) | 12,90059 | 5,14 | 8,396442 | 4,006969 |

\* Weiss and Wardrop [18], \*\* Schmidt et al. [19], 'Macari and Luqueti [20],” Borsa, Borsa and Borsa. [21].
RV = Reference Values; Globular volume = GV; number of red blood cells = Heme; mean corpuscular volume = MCV; total plasma protein = TPP; ns = not significant.
1 = linear regression ( $p = 0.0225$ ), Heme = $1686824 + 1091.2\text{DEB}$ , $R^2 = 0.73$ ;
2 = linear regression ( $p = 0.0351$ ), MCV = $164.9772796 - 0.0572823\text{DEB}$ , $R^2 = 0.57$ .

**Table 6.** Levels of dietary electrolyte balance on the leucogram of broilers from 01 to 21 days of age.

| DEB(mEq/Kg) | Leu ($\mu\text{L}$) | Het ($\mu\text{L}$) | Lyn ($\mu\text{L}$) | Eos ($\mu\text{L}$) | Bas ($\mu\text{L}$) | Mon ($\mu\text{L}$) | H/L |
| --- | --- | --- | --- | --- | --- | --- | --- |
| 110 | 18286 | 5977 | 10193 | 47,1 | 0,0 | 2524,0 | 0,6067 |
| 175 | 20833 | 6797 | 12737 | 95,7 | 0,0 | 2941,4 | 0,5580 |
| 240 | 19429 | 6941 | 10186 | 0,0 | 40,0 | 2498,3 | 0,6756 |
| 305 | 16833 | 5909 | 8525 | 0,0 | 74,3 | 2498,3 | 0,6319 |
| 370 | 18000 | 6580 | 9297 | 20,0 | 0,0 | 2102,9 | 0,7726 |
| VR | 12,000 – 30,000* | 3000 – 6000* | 7000 – 17,500* | 0 – 1000* | raro* | 150 – 2000* | 0,5** |
| Probability | 0,7265 | 0,9346 | 0,1870 | 0,2144 | 0,3323 | 0,5795 | 0.0532 |
| Regression | Ns | Ns | Ns | Ns | Ns | ns | Ns |
| CV | 28,12 | 43,11 | 31,19 | 262,98 | 355,04 | 36,56 | 26.62 |

\* Weiss and Wardrop [18], \*\* Macari and Luqueti [20].
RV = Reference Values; Leukocyte number = Leu; heterophiles = Het; lymphocytes = Lyn; eosinophils = Eos; basophils = Bas and monocytes = Mon; heterophile / lymphocyte ratio = H / L; ns = not significant.

**Table 7.**
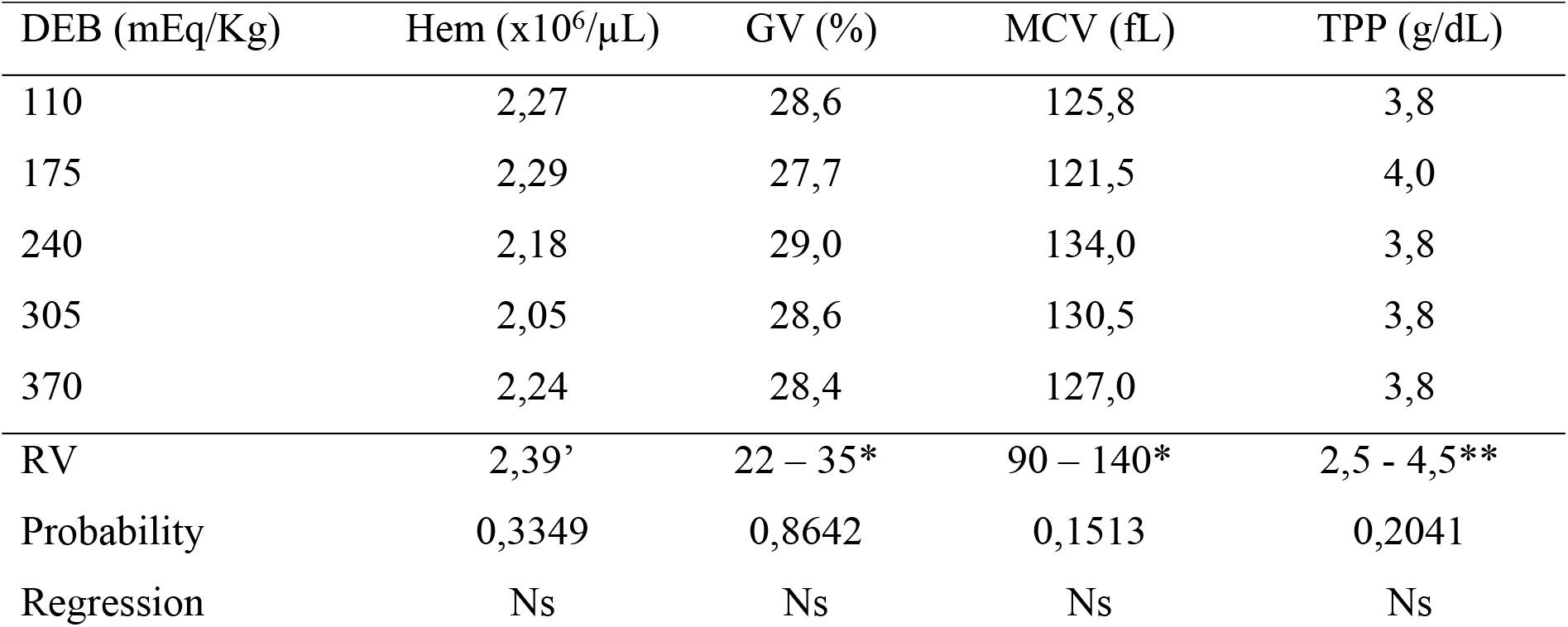
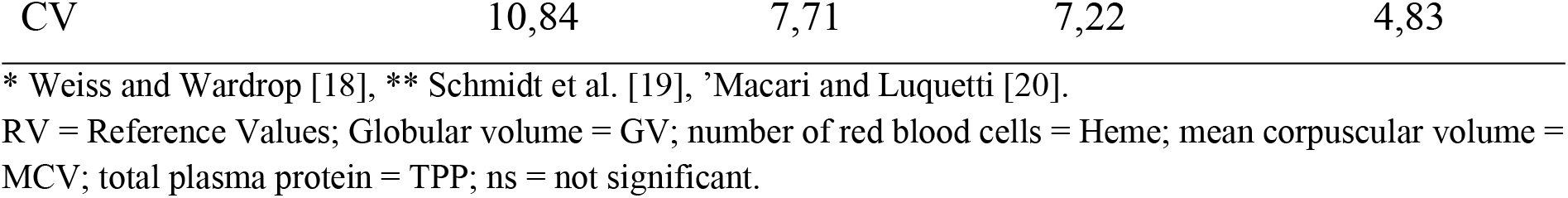
Levels of dietary electrolyte balance on the erythrogram and TPP of broilers from 22 to 42 days of age.

**Table 8.** Levels of dietary electrolyte balance on the leukogram of broilers at the 22 to 42 day old phase.

| DEB(mEq/Kg) | Leu (μL) | Het (μL) | Lyn (μL) | Eos (μL) | Bas (μL) | Mon(μL) | H/L |
| --- | --- | --- | --- | --- | --- | --- | --- |
| 110 | 20571 | 10316,7 | 7683 | 85,7 | 31,4 | 1956,0 | 1,3117 |
| 175 | 17143 | 6603,3 | 7460 | 0,0 | 25,7 | 2220,0 | 0,9635 |
| 240 | 15714 | 7241,4 | 6269 | 47,1 | 0,0 | 1922,0 | 1,0980 |
| 305 | 15143 | 5918,0 | 7153 | 32,9 | 22,9 | 1820,0 | 0,8383 |
| 370 | 17571 | 7210,0 | 7806 | 100,0 | 51,4 | 2096,0 | 0,9413 |
| VR | 12,000 – 30,000* | 3000 – 6000* | 7000 – 17,500* | 0 – 1000* | raro* | 150 – 2000* | 0,5** |
| Probability | 0,1043 | 0,0005 | 0,6899 | 0,5899 | 0.8396 | 0.8625 | 0.0532 |
| Regression | Q<sup>1</sup> | Q<sup>2</sup> | Ns | Ns | ns | Ns | L<sup>3</sup> |
| CV | 22,39 | 20,16 | 29,69 | 238,28 | 312,39 | 34.43 | 26.62 |

\* Weiss and Wardrop [18], \*\* Macari and Luqueti [20].
RV = Reference Values; Leukocyte number = Leu; heterophiles = Het; lymphocytes = Lyn; eosinophils = Eos; basophils = Bas and monocytes = Mon; heterophile / lymphocyte ratio = H / L; ns = not significant.
Averages with the same capital letters in the same column do not differ statistically according to the SNK test (p <0.05)
1 = quadratic regression (p = 0.0237), Leu = 30628 - 114.32DEB + 0.213DEB<sup>2</sup>, R<sup>2</sup> = 0.98;
2 = quadratic regression (p = 0.0029), Het = 16427 - 73.053DEB + 0.130DEB<sup>2</sup>, R<sup>2</sup> = 0.82.
3 = linear regression (p = 0.0238), H / L = 1.343438462 - 0.001293077DEB, R<sup>2</sup> = 0.164008

For the 22 - 42 days old phase, the Leu and Het variables had a quadratic effect (p <0.05) for the levels of electrolyte balance. The levels of 268 and 280 mEq of DEB/kg of feed were estimated by the derivative of the quadratic equations, which provided lower amounts of Leu and Het, respectively. There was no effect (p> 0.05) on the variables GV, Heme, MCV, TPP, Lyn, Eos, Bas and Mon depending on the tested DEB levels. A value above the reference of the species was observed for variable Het at all levels tested, except for the level 305 mEq / kg. At level 175 and 370 mEq/kg a value was observed above the reference for Mon and the H/L of all levels was also high, for the level of 240 mEq/kg the number of Lyn is below the reference value for the species. The average maximum and minimum temperature during this phase was 34.8 and 21.4°C, respectively, and the maximum and minimum humidity of 65.10 and 28.24%, and for the day of collection the maximum and minimum temperature was respectively 35,8 and 23.8 °C, and the maximum and minimum humidity 58 and 29%, characterizing chronic stress due to cyclic heat.

### Biochemistry

The results of the biochemical parameters are presented in Table 9 and 10 for the phase of 1 - 21 and 22 - 42 days of age, respectively. For the 1-21 days old phase, the variable chlorine (Cl) showed a linear effect (p <0.05) for the levels of electrolyte balance tested. There was no effect (p> 0.05) on the variables calcium (Ca), phosphorus (P), uric acid (UA) and urea (U) depending on the tested DEB levels. It was observed a value below the species reference for the variable P on all levels tested.

**Table 9.**
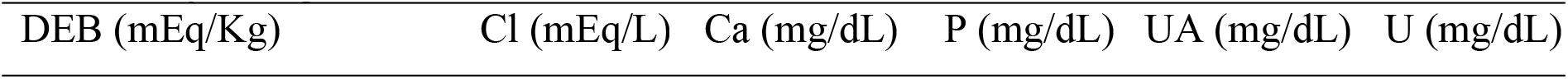
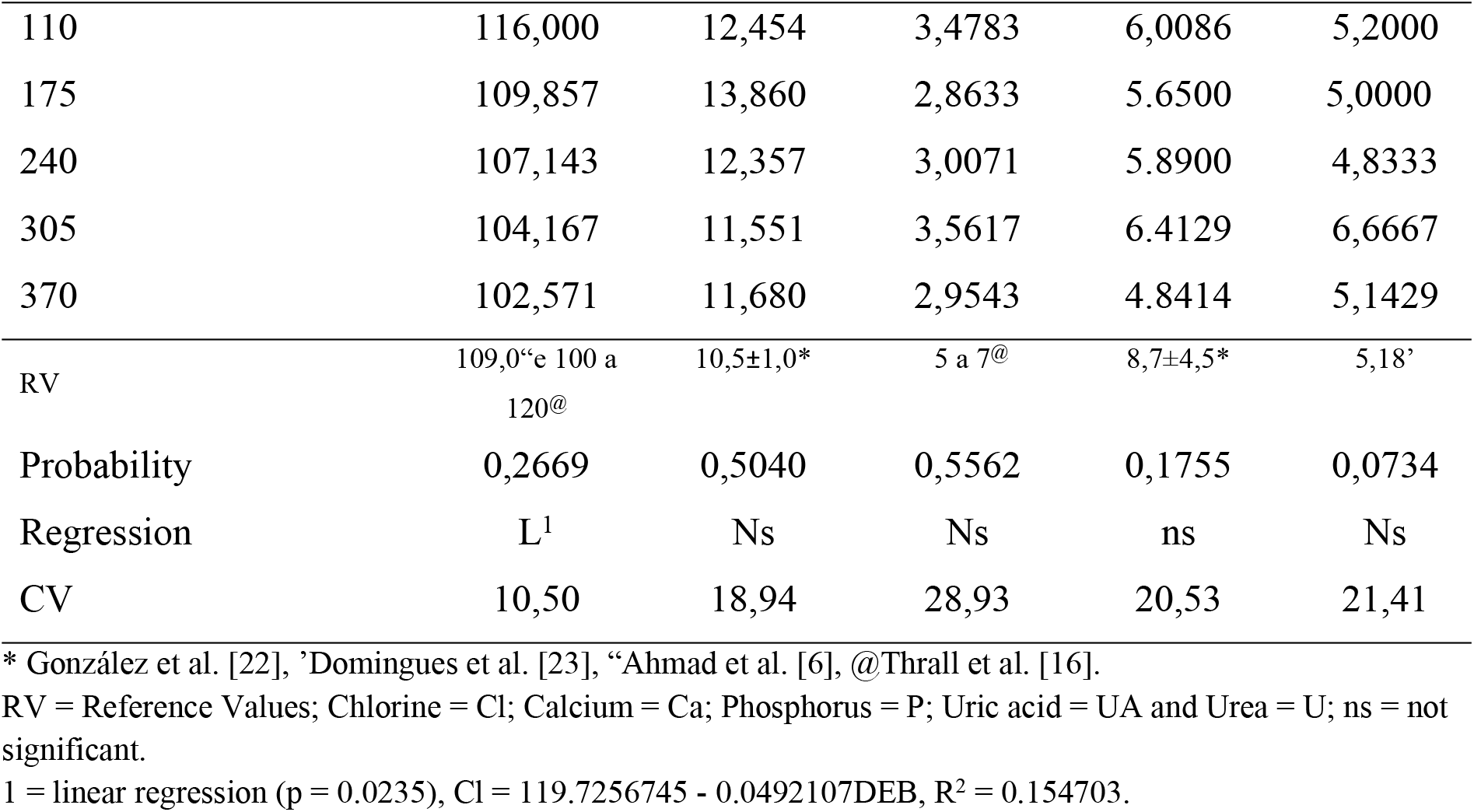
Levels of dietary electrolyte balance on serum biochemical parameters of broilers from 01 to 21 days of age.

**Table 10.**
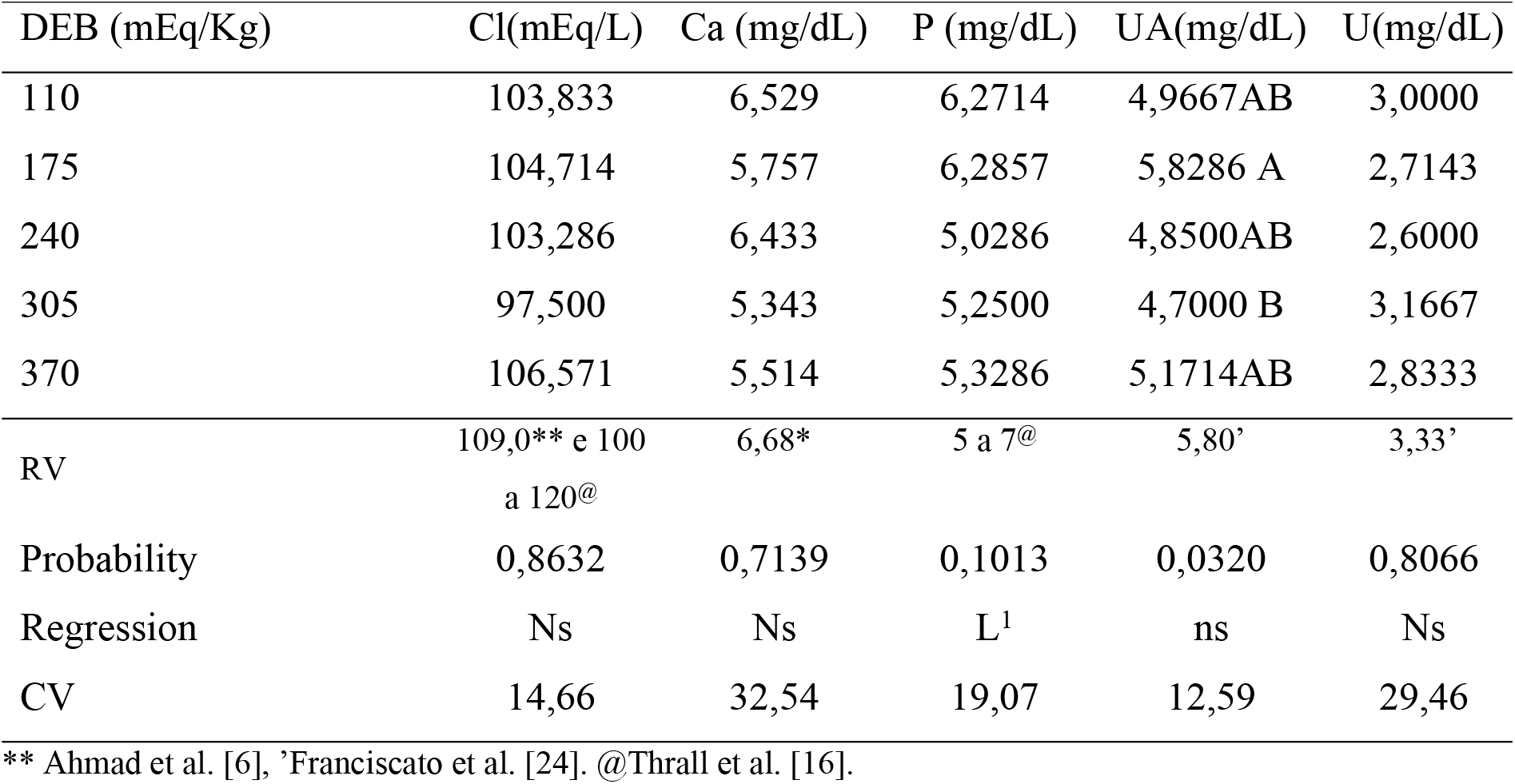
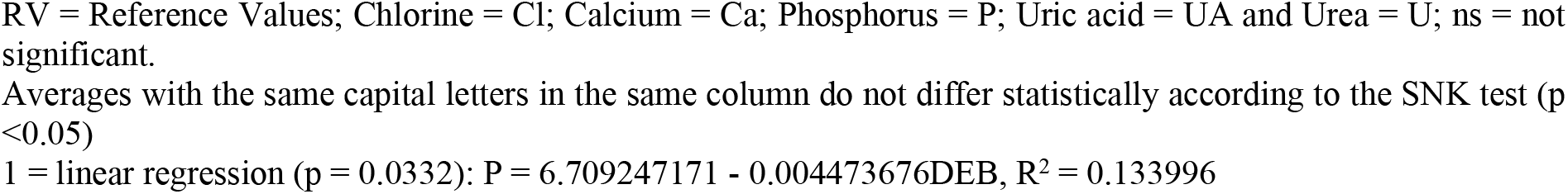
Levels of dietary electrolyte balance on the serum biochemical parameters of broilers from 22 to 42 days of age.

| DEB (mEq/Kg) | Cl(mEq/L) | Ca (mg/dL) | P (mg/dL) | UA(mg/dL) | U(mg/dL) |
| --- | --- | --- | --- | --- | --- |
| 110 | 103,833 | 6,529 | 6,2714 | 4,9667AB | 3,0000 |
| 175 | 104,714 | 5,757 | 6,2857 | 5,8286 A | 2,7143 |
| 240 | 103,286 | 6,433 | 5,0286 | 4,8500AB | 2,6000 |
| 305 | 97,500 | 5,343 | 5,2500 | 4,7000 B | 3,1667 |
| 370 | 106,571 | 5,514 | 5,3286 | 5,1714AB | 2,8333 |
| RV | 109,0** e 100<br>a 120<sup>@</sup> | 6,68* | 5 a 7<sup>@</sup> | 5,80' | 3,33' |
| Probability | 0,8632 | 0,7139 | 0,1013 | 0,0320 | 0,8066 |
| Regression | Ns | Ns | L<sup>1</sup> | ns | Ns |
| CV | 14,66 | 32,54 | 19,07 | 12,59 | 29,46 |

\*\* Ahmad et al. [6], 'Franciscato et al. [24]. @Thrall et al. [16].
RV = Reference Values; Chlorine = Cl; Calcium = Ca; Phosphorus = P; Uric acid = UA and Urea = U; ns = not significant.
Averages with the same capital letters in the same column do not differ statistically according to the SNK test ( $p < 0.05$ )
l = linear regression ( $p = 0.0332$ ): $P = 6.709247171 - 0.004473676\text{DEB}$ , $R^2 = 0.133996$

For the phase of 22-42 days of age, the variable P showed a linear effect (p <0.05) in order of the DEB levels tested, there was also a significant difference (p <0.05) between the means UA variable only between the levels of 175 and 305 mEq of DEB/Kg of feed. There was no effect (p <0.05) on the variables Cl, Ca and U due to the tested DEB levels. A value below the species reference was observed for the variable Cl at the level 305 mEq/Kg.

## Discussion

Electrolyte supplementation is used to mitigate the harmful effects of heat stress, in order to balance electrolyte losses by birds in thermal stress [6,7,8]. Quantitative and morphological changes in blood cells are associated with infectious, inflammatory conditions and stresses of a diverse nature, as we can see by variations in hematocrit values, number of circulating leukocytes and erythrocytes and hemoglobin content in red blood cells [25].

In a situation of chronic thermal stress, it is expected that there will be a decrease in the total number of red blood cells associated with a shorter lifespan and inhibition of production by the bone marrow, a decrease in MCV and a decrease in the number of leukocytes associated with lower feed consumption and an increase in costicosterone that depresses the activity of lymphoid organs, and a lower level of TPP [26, 13].

In previous studies it has been observed that the use of DEB for broilers improves the performance of birds, body weight gain, bone mineralization, decreased mortality and the normalization of the acid-basic balance, making softer the effect of thermal stress keeping hematological and biochemical parameters within the normal range. [10, 9, 6].

In this study, it was observed that in the phase of 1 - 21 days of age there was a linear effect (p <0.05) for the parameters, Heme and MCV, where an increase was observed proportionally to the DEB levels of Heme and an inversely proportional decrease of the MCV, both within the reference value for species, suggesting for this stage that the red blood cells have a longer useful life proportional to the increase in the level of DEB, considering that the red blood cells were increasing in number and decreasing in size. This can be considered, since according to Christian [27], how higher the animal’s metabolic rate, bigger the accumulated oxidative stress and the lower the average life of the erythrocytes. What reinforces this observation is that in addition to be increased in number, it was decreasing in size, indicating that it was living longer, even in the senescence process, as observed by Bartosz [28] the older the red blood cell, the smaller its size.

There was also a linear effect (p <0.05) for the level of Cl, in which its level decreases as the DEB increased. It has already been observed that thermal stress decreases the serum level of Cl- [13, 10], electrolyte is necessary in body fluids to exert an acidic action to normalize blood pH [11]. Ahmad et al. [6], also observed a linear decrease (p <0.05) in Cl as the DEB increased. In this study, Cl was within the reference value for the species in all DEB levels tested, however the linear decrease in its level indicates that values above the level of 370 would cause a state of hypochloremia, and it is not recommended to use values above this.

Monocytosis and hypophosphatemia were common to all DEB levels tested for this same phase. Monocytosis in birds is common in cases of infectious diseases caused by microorganisms that promote granulomatous inflammation such as Mycobacterium, Chlamydia and fungi, chronic bacterial granuloma and extensive tissue necrosis and zinc deficiency [16], while hypophosphatemia is common in the growth phase of young birds, hypovitaminosis D3 (with hypocalcaemia), malabsorption, starvation and prolonged corticosteroid therapies [16].

No evidence has been identified to justify these two situations then further studies need to be carried out. As the reference values are not always suitable for all populations, these are probably normal values for animals in this production phase, under local environmental conditions. Likewise, the heterophilia observed at levels 175, 240 and 370 mEq/kg, is questionable, as in studies already carried out in the same region, mean values of around 6412 μL have already been observed for broilers aged around 21 days [29], similar to that observed at this stage.

In this study, it was observed that in the 22 - 42 days of age of the birds, there was a quadratic effect (p <0.05) for the Leu and Het variables and a linear effect for the H / L variable as a function of the DEB levels tested. There was no variation in the erythrocyte parameters, in the total plasma proteins and in the number of lymphocytes, eosinophils, basophils and monocytes for this phase for the levels of DEB tested, similar to the one observed by Laganá et al. [30] and Xie et al. [12]. A value above the species reference for Het was observed in all levels tested except for the level 305 mEq/kg, however this heterophilia is questionable because, in works already carried out with broilers aged around 42 days in the same region, average values for this variable around 7020 μL were observed [29], similar to what was observed in this study for this phase, excepting for 110 mEq/kg level, where we observed a Het level above this average, therefore presenting heterophilia.

For all DEB levels the H/L ratio was above what is considered ideal for species, according to Gross and Siegel [31], there are three values based on the H/L ratio that can be used to characterize stress levels in broilers. In this classification, the value 0.2 would indicate a mild degree of stress; 0.5 intermediate or moderate stress and 0.8 high stress, however Macari and Luquetti [20] consider 0.5 the ideal proportion of H/L, and Laganá et al. [30] observed mean values of 0.79 in a thermoneutral environment (21-25 ° C) for H/L.

Considering this classification and Macari and Luquetti [20] and Laganá et al. [30] for all levels of DEB tested, the animals were under high stress, however the quadratic effect on Leu and Het demonstrates an improvement in leukocyte parameters as it approaches the 305mEq/kg DEB level. Both, the heterophilia at the 110 mEq/kg DEB level and the lymphopenia present at the 240 mEq/kg level, are associated with the high stress level that these animals were experiencing, there was an unusual variation in Mon (monocytosis) for the 175 and 370 mEq/kg level, and no other similar situation was observed in any other study.

The P variable presented a linear effect (p <0.05) for the DEB levels tested, different from what was observed by Vieites et al. [32], who found no variation for the levels of 0 to 350 mEq of DEB/kg on phosphorus. Junqueira et al. [33] found changes in P levels in diets with different DEB and reported that dietary sodium, when absorbed, combines with phosphorus plasma, forming sodium phosphate, and therefore facilitates its elimination by the kidneys, it has beneficial effects on the incorporation of calcium into the carbonate ion, with the consequent increase in calcium carbonate synthesis, what may justify the decrease in the serum phosphorus level as the DEB level increases in this experiment. There was also a significant difference (p <0.05) between the average of the UA variable only between the level 175 and 305 mEq of DEB/Kg, no relationship was identified for this decrease, however the levels remained within the reference values for the species.

The Hypochloremia, at the level of 305 mEq DEB/kg, needs to be further investigated, as according to Thrall, et al. [16] it is a rare phenomenon in birds and a similar situation has not been identified in other studies for DEB levels close to this.

On the 22 - 42 days old phase, the level of 268 to 280 mEq of DEB/kg of feed, showed a better response from the broilers immune system, because, within this range, the Leu and Het numbers were lower than in the others DEB levels tested and within the normal range for the specie. These values were obtained using quadratic regression equations.

## Conclusion

Supplementation with 370 mEq DEB/Kg of feed improves the hematological and biochemical response, it provided an increase in the number of red blood cells, without promoting metabolic imbalance in the phase from 1 until 21 days old. On the phase from 22 until 42 days old, the levels of 268 to 280 mEq of DEB/kg of additional feed improve the response from the broilers immune system.

